# Lévy Flight Patterns in the Cortical Architecture of *Macaca mulatta*

**DOI:** 10.1101/2025.01.29.635444

**Authors:** Arturo Tozzi

## Abstract

Lévy flights (LF), a concept originating in statistical physics, describe random walks in which the step lengths follow a heavy-tailed probability distribution, often a power law. Unlike Brownian motion, where step lengths are constrained within a narrow range, LF are characterized by the coexistence of many short steps interspersed with occasional long jumps. Applying advanced computational techniques, we looked for LF-like patterns in high-resolution histological images of *Macaca mulatta* (Rhesus macaque) cortical area 4 from BrainMaps.org. Step-length distributions, derived from pairwise distances between neuronal somata, exhibited heavy-tailed behavior consistent with power-law models across all samples. Maximum likelihood estimation of power-law exponents (α values: 0.87-1.08) strongly supported the heavy-tailed nature of these patterns, showing a better fit with power-law models compared to exponential or normal distributions. Connectivity analyses revealed a dual organizational structure within cortical layers: densely interconnected local clusters coexisting with sparse long-range connections. k-Nearest neighbors graphs demonstrated small-world network properties, with average clustering coefficients ranging from 0.622 to 0.630 across samples. This consistent structural organization aligns with LF principles, wherein local processing is optimized alongside global integration for efficiency and functionality. The implications extend to developmental biology, as the emergence of LF-like patterns likely reflects intrinsic self-organizing processes during embryonic and fetal development. This LF-like organization provides a natural framework for designing artificial networks that optimize performance in tasks requiring both localized specialization and global integration. Moreover, understanding the developmental origins of these patterns could guide strategies for neural repair and regeneration in stroke or neurodegenerative diseases.

## INTRODUCTION

The cortex of *Macaca mulatta* (Rhesus macaque) serves as an excellent model for studying laminar organization and heterogeneous connectivity, thanks to its evolutionary closeness to humans and the intricate complexity of its cortical structure (Distler and Hoffmann, 2015; Testard et al., 2024). Within this context, we explore a novel perspective on neuronal spatial organization: the potential existence of Lévy flight patterns. Lévy flights, a concept originating in statistical physics, describe random walks in which the step lengths follow a heavy-tailed probability distribution, often a power law (Boyer and Pineda, 2016). Unlike Brownian motion, where step lengths are constrained within a narrow range, Lévy flights are characterized by the coexistence of many short steps interspersed with occasional long jumps (Sims et al., 2012). This pattern has been observed in diverse natural systems, from animal foraging strategies and turbulent fluid flows to stock market dynamics and information-seeking behaviors. In biological systems, Lévy flights represent a mechanism for optimizing exploration and resource distribution, balancing local and global interactions (Campeau et al., 2022; Zhou et al., 2022; Papanikolaou et al., 2024). In the context of neuronal networks, this behavior could suggest an optimized balance between localized processing and long-range integration (Ohta et al., 2022).

In this study, we analyzed high-resolution histological images of cortical area 4 from *Macaca mulatta* to investigate the presence of Lévy flight-like like organization in the neuronal distributions. We compared multiple samples from BrainMaps.org, which provides interactive, multiresolution brain atlases of various animals comprising over 140 million megapixels. Previous studies have demonstrated that neurons in the cortex are not randomly distributed; rather, they exhibit clustering tendencies driven by developmental, functional and evolutionary constraints (Manley et al., 2024). These clusters often represent functional microcircuits, such as cortical columns or minicolumns, which facilitate local processing (Hosoya, 2019). However, neuronal connectivity is not limited to local interactions. Long-range connections are crucial for integrating information across distant regions of the cortex, forming the basis of complex cognitive and sensorimotor behaviors (Fisher et al., 2024). The coexistence of local clustering and long-range connections raises the question of whether the spatial organization of neurons reflects principles consistent with Lévy flights.

Recent advances in computational neuroscience and network science have provided powerful tools to investigate the spatial organization of neuronal networks. Quantitative analyses of step length distribution, power-law fitting, network-based clustering and connectivity patterns enable the identification of statistical signatures of Lévy flights (Tsubo et al., 2012; Tring et al., 2023; Armonaite et al., 2024). Specifically, step lengths between neuronal somata can be analyzed to determine whether they follow a heavy-tailed distribution, indicative of Lévy flight behavior. Additionally, graph-based connectivity analyses provide complementary insights into local and global network organization, highlighting clustering tendencies and long-range interactions. Together, these methods offer a robust framework for exploring the spatial organization of cortical neurons.

The presence of Lévy flight patterns in cortical layers would have profound implications for understanding the principles underlying cortical organization. Lévy flights are inherently efficient, enabling systems to balance exploration and exploitation. In the context of the cortex, this could reflect an optimal trade-off between local processing within clusters and global integration across cortical regions. This pattern would be consistent with the “small-world” topology observed in many biological networks, where dense local clustering coexists with sparse long-range connections (Suo eta al., 20217; Luboeinski et al., 2023; Bowen et al., 2024).

The paper is organized as follows. In the next section, we detail the methods used for image processing, point extraction and statistical analysis. We then present the results of the step length distribution analysis, power-law fitting and connectivity analyses. Finally, we discuss the implications of these findings in the context of cortical organization, focusing on the developmental and evolutionary processes shaping neuronal architecture, their impact on network efficiency and the broader relevance of Lévy flights in biological systems

## MATERIALS AND METHODS

### Samples and data preprocessing

We utilized high-resolution histological images of cortical area 4 from the BrainMaps.org database, which provides sub-micron resolution data (**Figure 1**). Three images from different individuals of *Macaca mulatta* were selected based on their high quality and representation of the cortical laminar structure. Image processing began with grayscale conversion to remove color information and standardize pixel intensity values. Noise reduction techniques, including Gaussian smoothing, were employed to reduce high-frequency noise that could interfere with segmentation and to minimize artifacts without compromising the integrity of the neuronal structures (Usman et al., 2021). The Gaussian kernel size was optimized through testing on images to ensure sufficient noise reduction while preserving neuronal boundaries. Contrast enhancement followed, utilizing histogram equalization to improve the dynamic range of pixel intensities and highlight the somata against the background (Ting et al., 2015).

**Figure 1.**
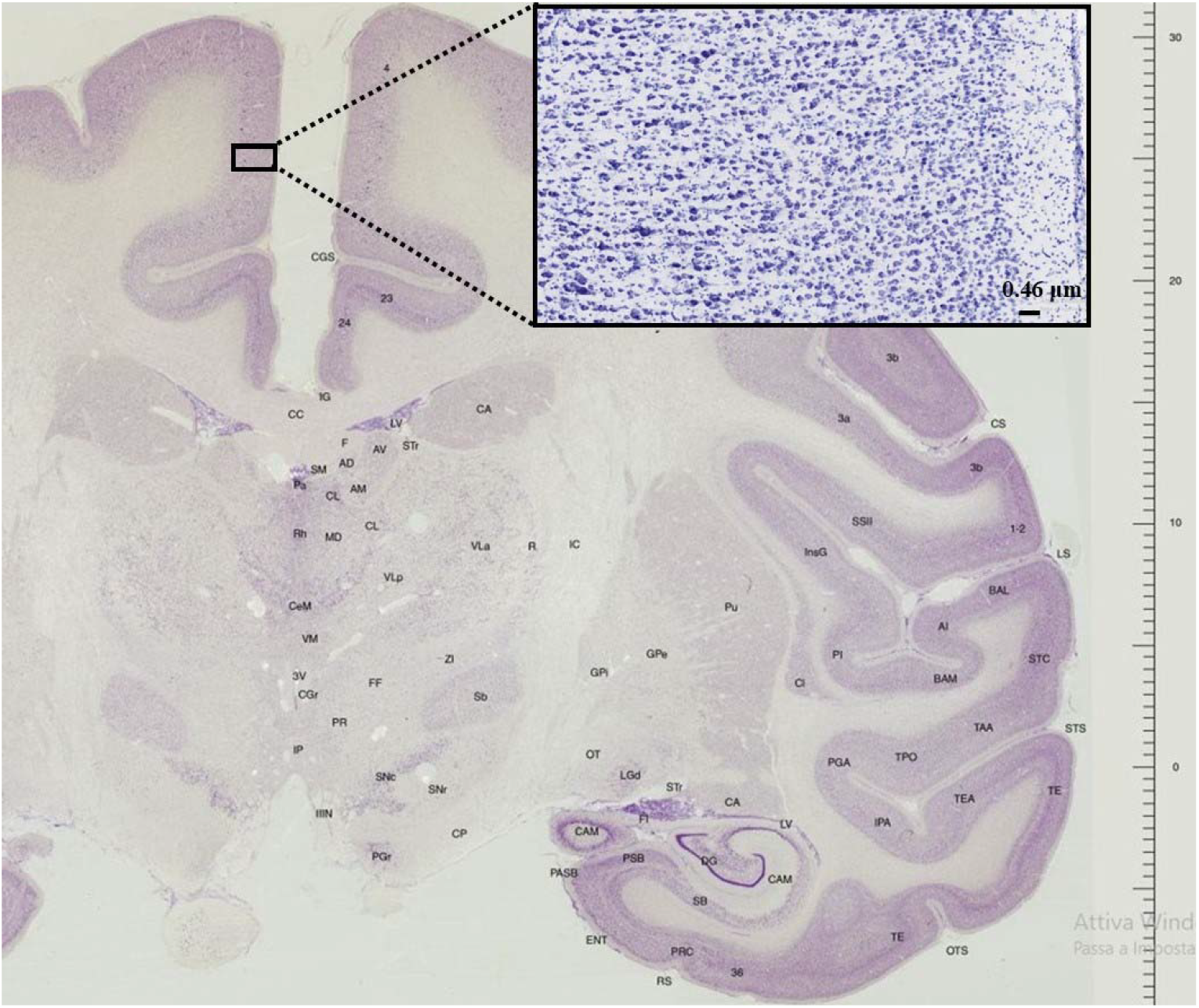
Cross-sectional view of the Macaca mulatta brain, with key reference points labeled. The inset provides a magnified view of the cortical layer within area 4, which was the focus of our analysis.

### Neuronal segmentation

Next, neuronal somata were identified using threshold-based segmentation, which relies on adaptive thresholding techniques to account for staining variability within and across samples (Shen et al., 2018). The darkest 20% of pixels in each image were isolated as regions of interest. This threshold was empirically chosen to maximize the inclusion of neuronal somata while minimizing background artifacts. Morphological operations, such as erosion and dilation, were employed to refine segmented regions (Huang et al., 2023). Erosion removed small, disconnected artifacts, while dilation reconnected fragmented somata slightly broken during thresholding. To avoid over-segmentation, a size filter excluded objects below a minimum area threshold, corresponding to the smallest known neuronal cross-sections, as determined from sample images.

### Step length distributions

Step lengths, defined as the pairwise Euclidean distances between neuronal somata, were computed to characterize the spatial relationships between neurons. These step lengths were used to generate step length distributions, which were evaluated for heavy-tailed behavior indicative of Lévy flights. Specifically, the distributions were plotted on logarithmic scales to visually assess their fit to a power-law model. Statistical fitting was performed using maximum likelihood estimation (MLE), which is robust to noise and small sample sizes (Milligan, 2003; Huang et al., 2015; Kwasniok, 2021). The power-law exponent (α) was estimated to quantify the heavy-tailed nature of the distribution, while goodness-of-fit tests were conducted to validate the model. To compare the step length distributions with alternative models, exponential and normal distributions were also fitted to the data. Parameters for these models were estimated using MLE and their fits were compared to the power-law model using likelihood ratio tests. This comparative analysis provided a statistical basis for determining whether the observed distributions were consistent with Lévy flight behavior.

### Connectivity analysis

In addition to step length analysis, the spatial organization of neurons was examined through connectivity analysis. A k-nearest neighbors (k-NN) graph was constructed for each sample, where nodes represented neuronal somata and edges connected each node to its k nearest neighbors based on Euclidean distance (Pei et al., 2023). The value of k was chosen to balance local and global connectivity, with sensitivity analyses performed to assess the robustness of the results to variations in k. The connectivity graph was analyzed to compute the average clustering coefficient, a measure of local interconnectedness (Ha et al., 2024). High clustering coefficients are indicative of dense local clustering, a hallmark of small-world networks. To visualize connectivity, graphs were plotted with nodes positioned according to their spatial coordinates, while edges were represented as lines connecting the nodes.

### Tools and statistics

Statistical comparisons were made using t-tests and bootstrapping techniques to estimate confidence intervals for the power-law exponent, clustering coefficient and exponential scale parameter across samples (Zahel 2022). All analyses were implemented in Python, leveraging libraries such as NumPy and SciPy for numerical computations, Matplotlib for visualization and NetworkX for graph-based analyses. Custom scripts were developed for image segmentation, point extraction and statistical modeling.

In summary, we investigated the potential coexistence of densely connected local clusters alongside long-range connections linking distant nodes. We assessed whether these features align with the hypothesized Lévy flight model, where short steps predominantly govern local connectivity, punctuated by occasional long jumps that enable global integration. If confirmed, these patterns would emphasize the structural organization of the neuronal network, illustrating the critical balance between local and global connectivity essential for efficient network function.

## RESULTS

The spatial organization of neurons in the cortical layers of *Macaca mulatta* was quantitatively analyzed across three histological samples (**Figure 2**). Step length distributions derived from the pairwise distances between neuronal somata revealed consistent patterns indicative of Lévy flight behavior across all samples. In each case, the distribution exhibited a heavy-tailed pattern, with a majority of short distances punctuated by occasional long jumps. When fitted to a power-law model using maximum likelihood estimation, the step length distributions demonstrated exponents (α) ranging between 0.87 and 1.08, confirming the presence of heavy-tailed behavior (**Figure 3**). Goodness-of-fit tests validated the robustness of the power-law model, which significantly outperformed exponential and normal distributions based on likelihood ratio tests.

**Figure 2.**
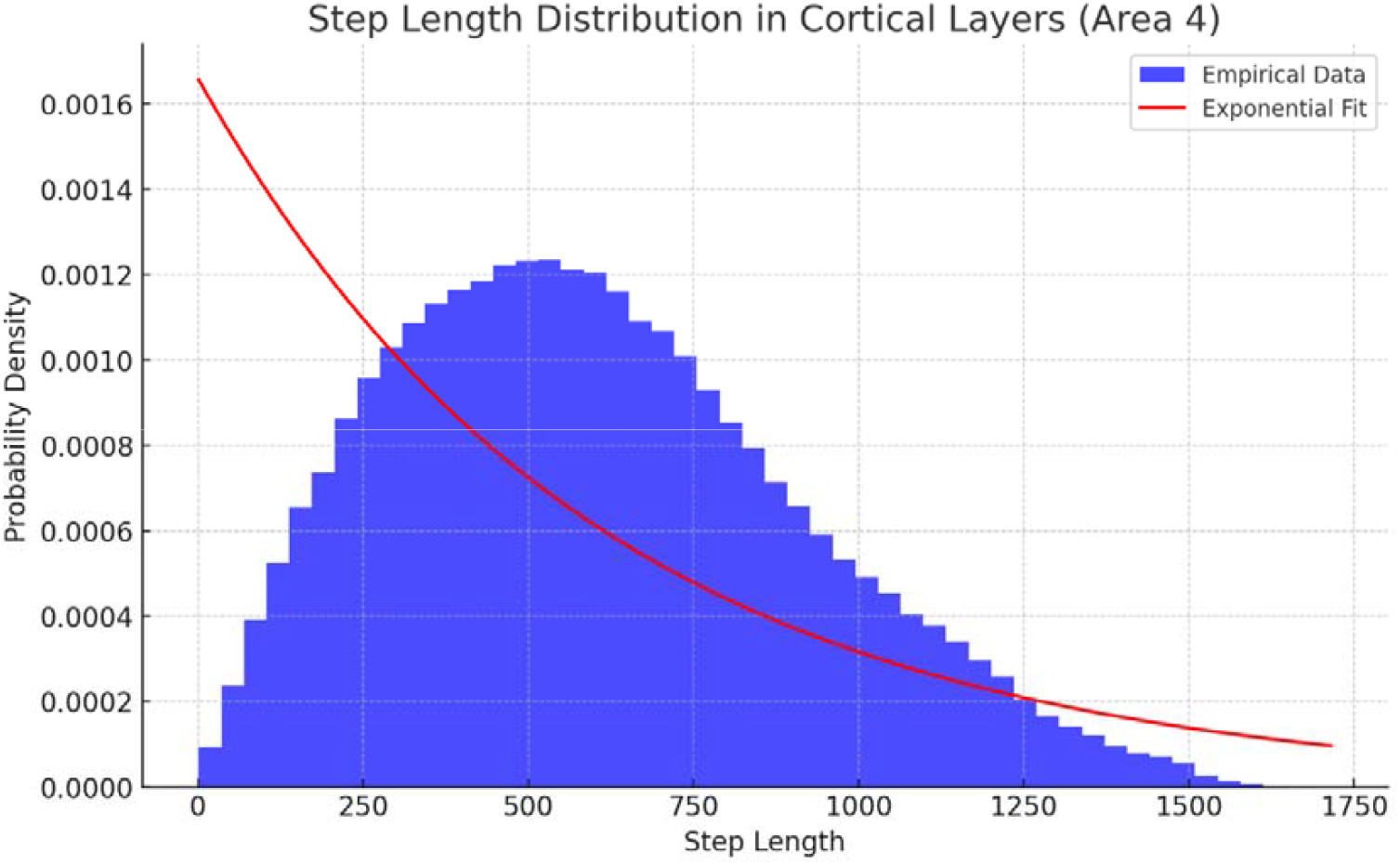
Distribution of step lengths, defined as the pairwise distances between neuronal somata, measured across the cortical layers of Area 4 in a single individual. The blue histogram displays empirical data, with a peak at shorter step lengths and a gradual decline at longer distances. The red curve, representing a fitted exponential distribution with a scale parameter of 603.48 (average step length), aligns well with shorter distances but significantly diverges in the tail. This divergence suggests heavy-tailed behaviour, characteristic of Lévy flights, which predict a greater likelihood of long distances compared to the exponential model.

**Figure 3.**
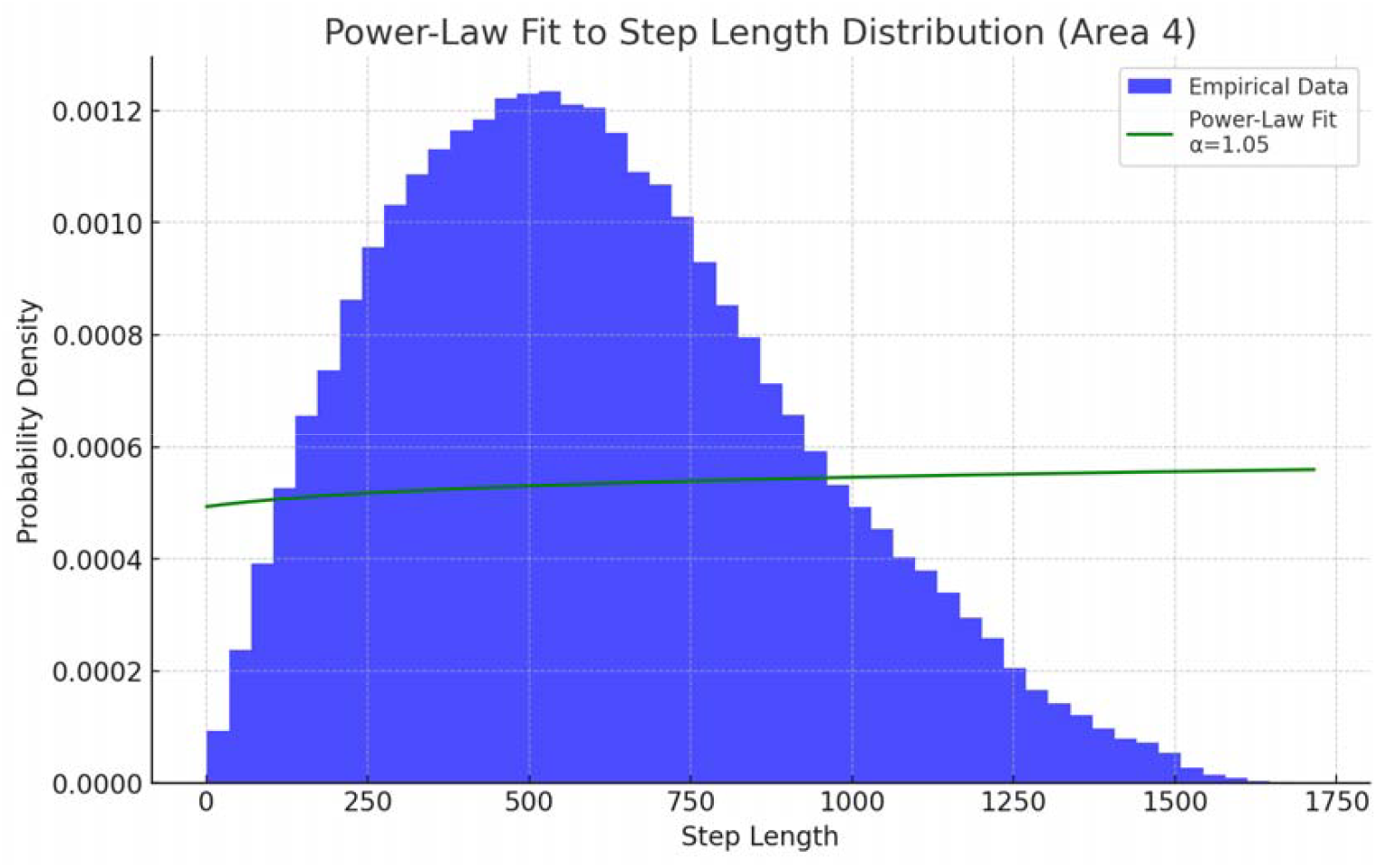
Power-law fit to the step length distribution, measured across the cortical layers of Area 4 in a single individual. The empirical step length distribution (blue histogram) shows predominantly short steps with a smaller proportion of long steps. The power-law fit (green curve), with a shape parameter α=1.05, indicates a heavy-tailed distribution, consistent with Lévy flight behaviour.

<u>**Figures 2 and 3** illustrate the step length distributions for two representative samples, overlaid with the fitted power-law and exponential models. The histograms reveal a steep decline in probability density for shorter step lengths, followed by a long tail representing ra</u>re but substantial distances. The power-law fit aligns closely with the empirical data in the tail regions, whereas the exponential model deviates substantially, failing to capture the heavy-tailed nature of the distribution. These findings provide evidence that the neuronal spatial organization in cortical area 4 adheres to principles consistent with Lévy flights.

The connectivity analysis further corroborated the presence of Lévy flight-like organization. k-Nearest neighbor graphs constructed from the spatial coordinates of neuronal somata revealed a dual structure: densely interconnected local clusters coexisting with sparse long-range edges (**Figure 4**). The average clustering coefficients, which quantify the extent of local connectivity, ranged from 0.622 to 0.630 across the three samples. These values are indicative of small-world network properties, where dense local clustering supports specialized processing and sparse global connections enable efficient integration across distant regions, pointing towards the coexistence of localized clusters and long-range connections.

**Figure 4.**
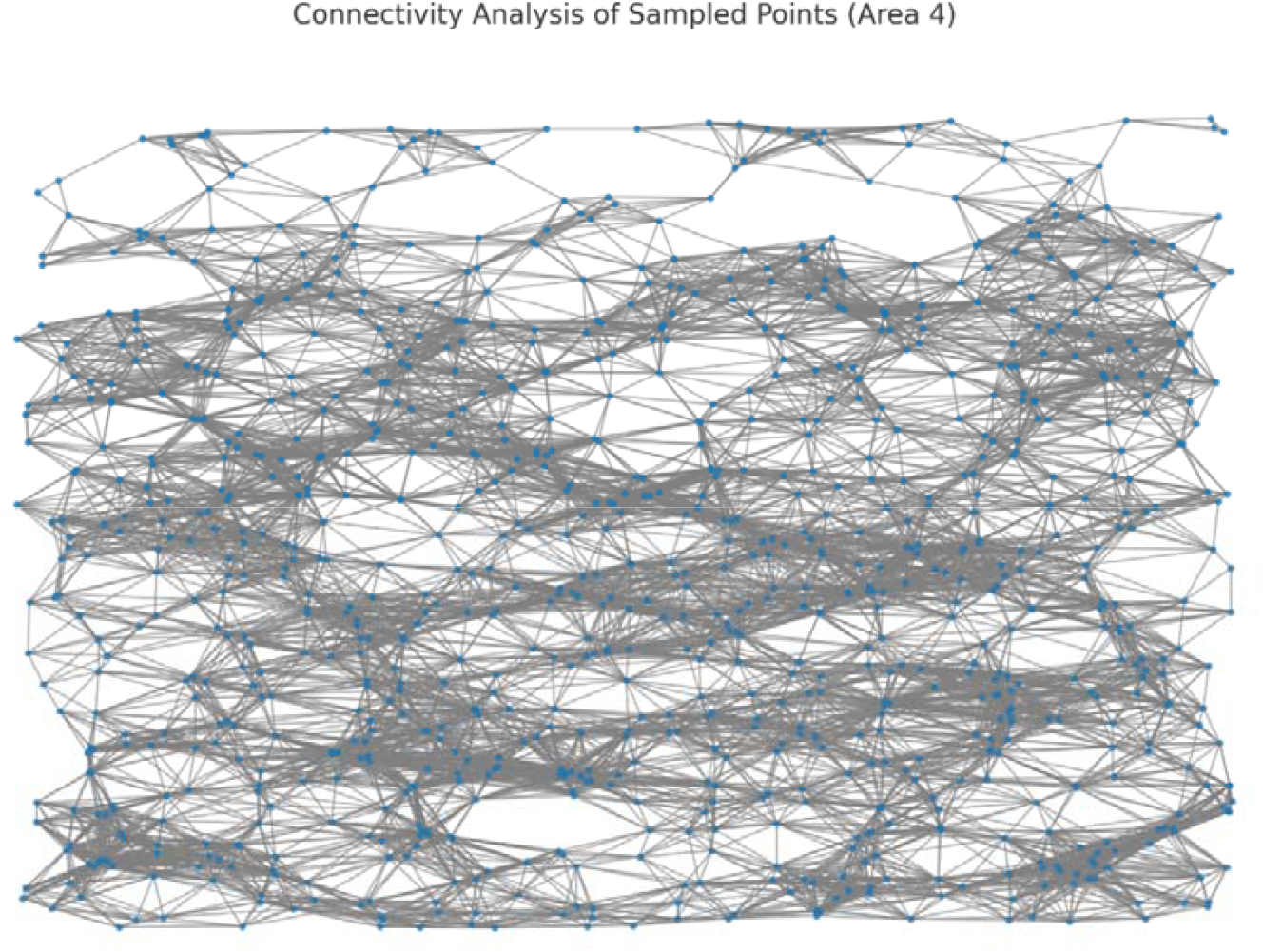
Connectivity analysis of sampled points within the cortical layers of Area 4 for a single individual. Blue dots represent nodes corresponding to sampled points of interest, such as darker regions in the image. Gray lines depict edges connecting nodes within a defined distance threshold, illustrating local connectivity between points. The graph reveals a combination of densely clustered regions and more sparsely connected areas

Comparative analyses across samples revealed minimal variability, suggesting that the observed patterns are intrinsic to the cortical layer rather than artifacts of the three samples (**Figure 5**). Pairwise t-tests conducted on the power-law exponents and clustering coefficients across samples yielded no statistically significant differences, underscoring the consistency of the findings. Bootstrapping further confirmed the robustness of the parameter estimates, with narrow confidence intervals for both the power-law exponents and clustering coefficients. These results indicate that the spatial organization of neurons in cortical area 4 is a conserved feature across individuals, reflecting fundamental principles of cortical architecture.

**Figure 5.**
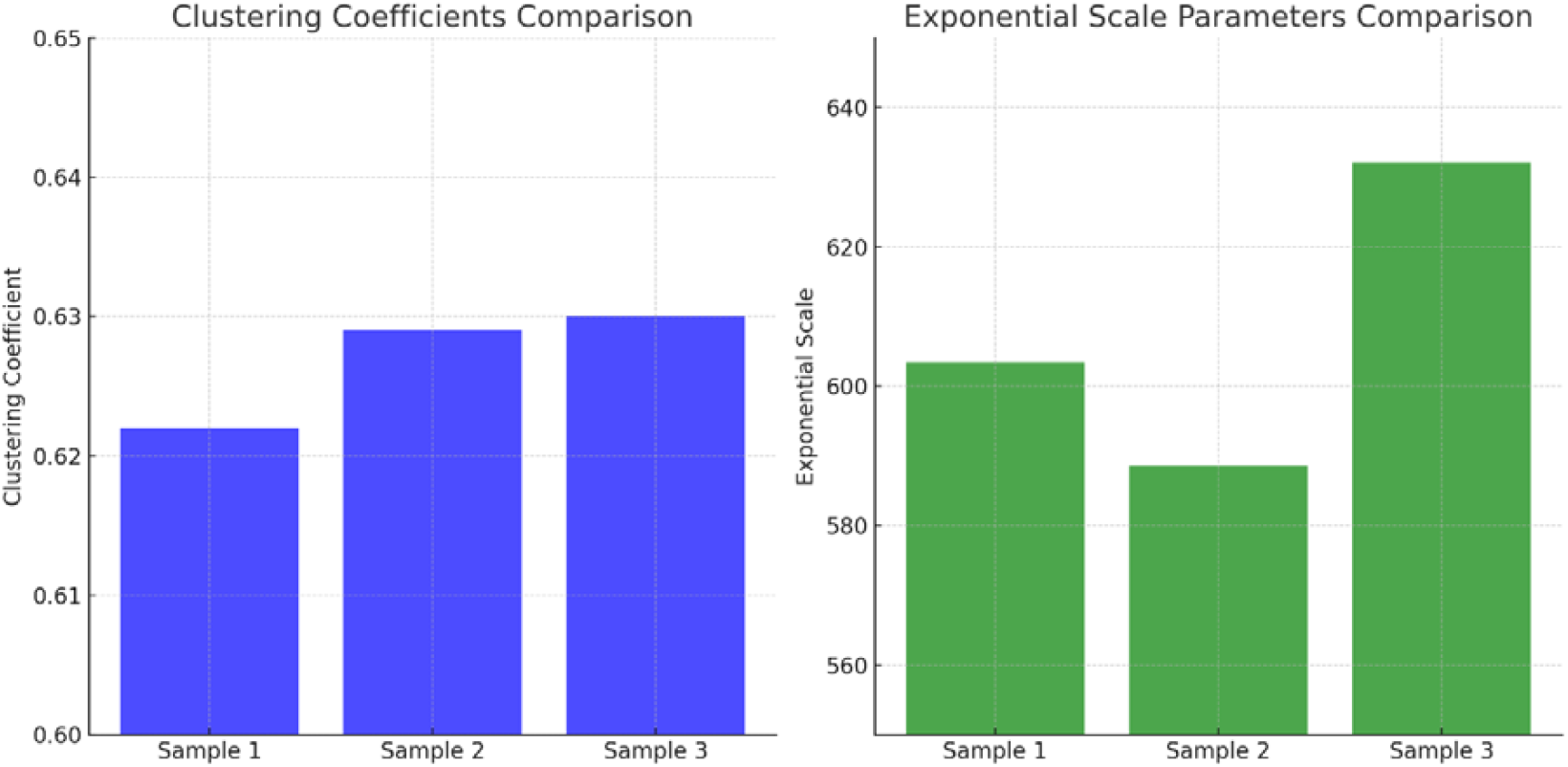
Comparison of exponential scale parameters and clustering coefficients across cortical layers of Area 4 in the three individuals. The clustering coefficients show minimal variation (0.622, 0.629, and 0.630), indicating consistently high local connectivity across samples, with t-test results being inconclusive due to near-identical values. Exponential scale parameters exhibit slight, nonsignificant variation (603.48, 588.49, and 632.10), reflecting natural biological variability rather than structural differences. These findings suggest that both clustering coefficients and step length distributions are conserved features of the cortical layers.

In addition to the quantitative metrics, visualizations of the connectivity graphs provided qualitative insights into the structural organization of the cortical layer. The graphs consistently displayed a balance between local and global connectivity, with densely connected nodes forming clusters interspersed with long-range edges linking distant regions. These features align with the hypothesized Lévy flight model, where short steps dominate but occasional long jumps enable global integration.

Overall, the results demonstrate that the spatial organization of neurons in cortical area 4 of *Macaca mulatta* is characterized by Lévy flight-like behavior. The heavy-tailed step length distributions, high clustering coefficients and connectivity graph structures collectively support this conclusion.

## CONCLUSIONS

We provide evidence that the spatial organization of neurons in the cortical layers of *Macaca mulatta* is characterized by Lévy flight-like behavior. This finding, based on quantitative analyses of step length distributions, power-law fitting and connectivity metrics, underscores the presence of a conserved, optimized spatial pattern in cortical area 4. The implications of these findings are far-reaching, extending from methodology to fundamental neuroscience and practical applications in developmental biology, computational modeling and brain-inspired systems.

One of the key advantages of this study lies in its methodological framework, which integrates high-resolution imaging with advanced statistical and computational techniques. By leveraging the details of BrainMaps.org, we were able to assess the neuronal spatial data at sub-micron resolution, ensuring precise and reliable measurements. The use of maximum likelihood estimation for power-law fitting provided robust parameter estimates, while connectivity analyses revealed structural patterns that are often obscured in conventional histological studies (Huang et al., 2015; Kwasniok, 2021; Pei et al., 2023). This integrative approach not only validated the presence of Lévy flight-like behavior but also highlighted its functional relevance. From an organizational perspective, the coexistence of densely clustered local connections with sparse long-range connections reflects an optimized balance between localized processing and global integration (Tozzi et al., 2018; Petersen et al., 2024). The duality between local clusters and long-range connections aligns with the small-world properties observed in many biological systems and provides a robust framework for understanding how the brain achieves both high efficiency and resilience (Luboeinski et al., 2023). The observation of consistent Lévy flight-like patterns across multiple samples suggests that these patterns are a conserved feature of cortical organization. This conservation implies that similar principles may be observed in other cortical regions or even across species. Comparative studies could explore whether Lévy flight-like behavior is a universal property of mammalian cortical organization or whether it varies depending on functional demands and evolutionary pressures.

One particularly intriguing avenue for future research lies in the relationship between Lévy flight-like patterns and embryonic and fetal development. The spatial organization of neurons in the cortical layers emerges during development through interplay of genetic, molecular and environmental factors (Arata et al., 2015; Caldarelli et al., 2024; Maniou et al., 2024). Early in development, neuronal migration and axonal growth are guided by gradients of signaling molecules such as netrins and semaphorins, that establish the initial layout of cortical circuits (Culotti and Kolodkin, 1996; Laclef and Métin, 2018; Zhang et al., 2028). The Lévy flight-like patterns may arise as an emergent property of these processes, reflecting an intrinsic tendency for systems to self-organize into efficient configurations (Zhang et al., 2000). Experimental predictions can be derived from this hypothesis, including the expectation that disruptions in signaling gradients during critical developmental windows may lead to deviations from Lévy flight-like spatial patterns, potentially impairing cortical functionality.

The implications of these findings extend beyond basic neuroscience to include practical applications in computational modeling and artificial intelligence. The Lévy flight-like organization observed in cortical layers provides a natural template for designing networks that balance efficiency and adaptability. Artificial neural networks inspired by this pattern could exhibit enhanced performance in tasks requiring both local specialization and global integration, such as image recognition or decision-making. Additionally, understanding the developmental processes that give rise to Lévy flight-like patterns may inform strategies for repairing or regenerating damaged neural tissue. By mimicking these developmental processes, it may be possible to guide the formation of functional networks in conditions such as stroke or neurodegenerative diseases.

Several limitations should be acknowledged. While the use of high-resolution histological images ensured precise measurements, the reliance on two-dimensional data introduces potential biases. The actual three-dimensional architecture of cortical layers may reveal additional insights into neuronal connectivity that are not captured in this study. Yet, the sample studied was too small to draw general conclusions. Further, while the power-law fitting and clustering analyses provided robust statistical evidence, the models used do not fully account for the influence of biological factors such as axonal guidance, synaptic plasticity and activity-dependent processes. These limitations highlight the need for complementary approaches, such as in vivo imaging and computational simulations, to further validate and extend the findings. Three-dimensional imaging modalities, such as serial block-face electron microscopy and light-sheet fluorescence microscopy, offer the potential to map cortical architecture in unprecedented detail (Smith and Starborg, 2019; Hobson et al., 2022; Marshall et al., 2023). Coupled with machine learning algorithms for image analysis, these techniques could automate the extraction of spatial data from large datasets. Furthermore, advancements in network science, such as the integration of multilayer network models, could provide deeper insights into the interplay between local and global connectivity in cortical layers.

In conclusion, the spatial organization of neurons in the cortical area 4 of *Macaca mulatta* exhibits Lévy flight-like behavior, characterized by heavy-tailed step length distributions, high clustering coefficients and small-world connectivity patterns. This conserved feature may reflect an optimized balance between local processing and global integration, supporting the efficient functioning of cortical networks. By exploring the developmental processes that shape these patterns, investigating their functional implications and harnessing them for practical applications, we can deepen our understanding of the principles governing cortical organization and their significance for brain health and disease. This study highlights the importance of cross-disciplinary collaboration in advancing our understanding of the brain cortex, by bridging concepts from statistical physics, developmental biology and neuroscience. Moving forward, collaboration between experimentalists, theorists and engineers will be essential for translating these insights into actionable knowledge.

## DECLARATIONS

### Ethics approval and consent to participate

This research does not contain any studies with human participants or animals performed by the Author.

### Consent for publication

The Author transfers all copyright ownership, in the event the work is published. The undersigned author warrants that the article is original, does not infringe on any copyright or other proprietary right of any third part, is not under consideration by another journal and has not been previously published.

### Availability of data and materials

all data and materials generated or analyzed during this study are included in the manuscript. The Author had full access to all the data in the study and take responsibility for the integrity of the data and the accuracy of the data analysis.

## Acknowledgements

none.

## Authors’ contributions

The Author performed: study concept and design, acquisition of data, analysis and interpretation of data, drafting of the manuscript, critical revision of the manuscript for important intellectual content, statistical analysis, obtained funding, administrative, technical and material support, study supervision.

## Declaration of generative AI and AI-assisted technologies in the writing process

During the preparation of this work, the author used ChatGPT to assist with data analysis and manuscript drafting. After using this tool, the author reviewed and edited the content as needed and takes full responsibility for the content of the publication.

